# AI-2 Production in *Fusobacterium nucleatum* Is Subspecies-Specific and Uncoupled from Quorum Sensing

**DOI:** 10.64898/2026.03.02.709096

**Authors:** G C Bibek, Shiqi Xu, Truman Tran, Chenggang Wu

## Abstract

Autoinducer-2 (AI-2) is a LuxS-dependent product of the activated methyl cycle (AMC) that functions as a quorum-sensing signal in diverse bacteria. *Fusobacterium nucleatum* is a genetically heterogeneous oral anaerobe comprising four subspecies: *nucleatum* (FNN), *vincentii* (FNV), *polymorphum* (FNP), and *animalis* (FNA). Previous studies have reported that FNN and FNP strains produce AI-2 and have proposed that AI-2–mediated quorum sensing contributes to biofilm formation and virulence. However, the distribution and functional relevance of AI-2 across all subspecies have not been systematically examined. Here, we show that AI-2 production is restricted to FNA strains. Genomic analysis revealed that FNN and FNV lack *luxS*, whereas FNP carries a disrupted *luxS* homolog. Consistent with these findings, AI-2 bioassays using the *Vibrio harveyi* BB170 reporter detected AI-2 exclusively in FNA strains. Deletion of *luxS* in FNA abolished AI-2 production but resulted in minimal transcriptional changes, and exogenous AI-2 failed to elicit global gene expression responses in non-producing subspecies. These results demonstrate that AI-2 production in *F. nucleatum* is subspecies-specific and uncoupled from quorum sensing. Our findings revise current assumptions regarding AI-2–mediated communication in *F. nucleatum* and reveal previously unrecognized metabolic divergence within the species complex.

**IMPORTANCE:** Periodontitis affects nearly half of adults in the United States and remains a leading cause of tooth loss worldwide. *Fusobacterium nucleatum* is a central member of oral biofilms and has also been linked to adverse pregnancy outcomes and colorectal cancer. Although AI-2–mediated quorum sensing has been proposed to contribute to its biofilm formation and virulence, our study demonstrates that AI-2 production is confined to subsp. *animalis* and is absent in other subspecies. Moreover, AI-2 does not function as a conserved quorum-sensing regulator in this species. These findings fundamentally revise prevailing assumptions about AI-2 signaling in *F. nucleatum* and suggest that subspecies-specific metabolic traits, rather than universal quorum sensing, may underlie ecological adaptation and host association.

## INTRODCUTION

Autoinducer-2 (AI-2) is widely regarded as a quorum-sensing signal that mediates communication within and between bacterial species and has been proposed to function as a “universal” language of microbial communities(1, 2). AI-2 is generated as a by-product of the activated methyl cycle (AMC), a central metabolic pathway that recycles S-adenosylmethionine (SAM), the universal methyl donor used for methylation of DNA, RNA, proteins, and metabolites(3–5). In the LuxS-dependent branch of the AMC, the enzyme LuxS cleaves S-ribosylhomocysteine to generate 4,5-dihydroxy-2,3-pentanedione (DPD), which spontaneously rearranges into AI-2. AI-2 accumulates extracellularly as bacterial cell density increases and, upon reaching a threshold concentration, is detected by specific receptors or transported into cells to regulate transcription in a density-dependent manner(6).

In the oral cavity, AI-2 has been implicated in the development of polymicrobial biofilms and in interspecies coordination(7). Multiple oral bacteria produce AI-2, including *Streptococcus mutans*, *Streptococcus gordonii*, *Streptococcus oralis*, *Aggregatibacter actinomycetemcomitans*, *Porphyromonas gingivalis*, and *Fusobacterium nucleatum*(8–13). Because of its central ecological role as a structural “bridge” organism linking early and late colonizers, *F. nucleatum* has received particular attention in studies of interspecies signaling(14).

*F. nucleatum* is a strictly anaerobic Gram-negative bacterium that plays a pivotal role in dental plaque formation through its broad coaggregation capacity (15, 16). Beyond its contribution to periodontal disease, it has been associated with adverse pregnancy outcomes and colorectal cancer following extraoral dissemination(17, 18). Importantly, *F. nucleatum* exhibits considerable genetic and phenotypic heterogeneity and is currently classified into four subspecies: subsp. *nucleatum* (FNN), *vincentii* (FNV), *polymorphum* (FNP), and *animalis* (FNA)(19–21). These subspecies differ in their ecological distributions and disease associations(22–24). FNN strains are frequently associated with periodontitis and exhibit strong biofilm-forming capacity, particularly in association with *Porphyromonas gingivalis*(22, 25–31). In contrast, FNA strains are more strongly associated with colorectal cancer and apical abscesses (23, 32), whereas FNP and FNV are more commonly detected in healthy oral sites (22, 29, 33). Despite these differences, the molecular mechanisms underlying subspecies-specific pathogenicity remain poorly understood.

Previous reports have suggested that *F. nucleatum* produces AI-2 and that AI-2–mediated quorum sensing contributes to biofilm formation and host interaction(7, 34–40). However, these studies focused on two individual strains (FNN ATCC 25586 and FNP ATCC 10953), and the distribution of AI-2 production across all four subspecies has not been systematically assessed. Moreover, whether AI-2 functions as a *bona fide* quorum-sensing signal in *F. nucleatum* rather than merely reflecting AMC activity remains unclear.

In this study, we investigated both the distribution and functional significance of AI-2 in *F. nucleatum*. Using the *Vibrio harveyi* BB170 reporter assay, we demonstrate that AI-2 production is restricted to the subsp. *animalis*. FNN, FNV, and FNP strains do not produce detectable AI-2 under the conditions tested. Furthermore, deletion of *luxS* in FNA abolished AI-2 production but caused minimal transcriptional changes, and exogenous AI-2 failed to induce global gene expression responses in non-producing subspecies. These findings indicate that AI-2 production in *F. nucleatum* is subspecies-specific and uncoupled from quorum sensing, fundamentally revising prevailing assumptions about AI-2–mediated communication in this species complex.

## RESULTS

### AI-2 Production Is Restricted to Subsp. animalis and Requires an Intact *luxS* Gene

Because AI-2 production depends on a functional *luxS* gene, we first examined the presence and integrity of *luxS* across the four subspecies of *F. nucleatum*. Using the LuxS protein from *P. gingivalis* ATCC 33277 (PGN_1474) as a query, we performed BLAST analyses against publicly available *F. nucleatum* genomes in the BioCyc and Integrated Microbial Genomes & Microbiomes (IMG/M) databases.

Unexpectedly, none of the analyzed FNN genomes contained a *luxS* homolog (Fig. 1A & 1B). This was surprising because previous studies reported that FNN strains—particularly the widely used model strain ATCC 25586—produce AI-2 (7, 34–39). Similarly, all examined FNV genomes also lacked *luxS*. In contrast, every FNA genome contained a single intact *luxS* gene, located between *uraA* (encoding an uracil-xanthine permease) and *pepF* (encoding oligoendopeptidase F) (Fig. 1A and 1B). FNP genomes also contained a *luxS*-like sequence at the same chromosomal position. However, this gene was interrupted by the insertion of a mobile genetic element belonging to the IS200 transposon family (Fig. 1B). The insertion disrupts the coding sequence near the 5′ end, preventing production of a full-length *luxS* mRNA and, consequently, a functional LuxS protein. This genomic data predicts that only FNA strains retain the capacity to produce AI-2.

**Figure 1.**
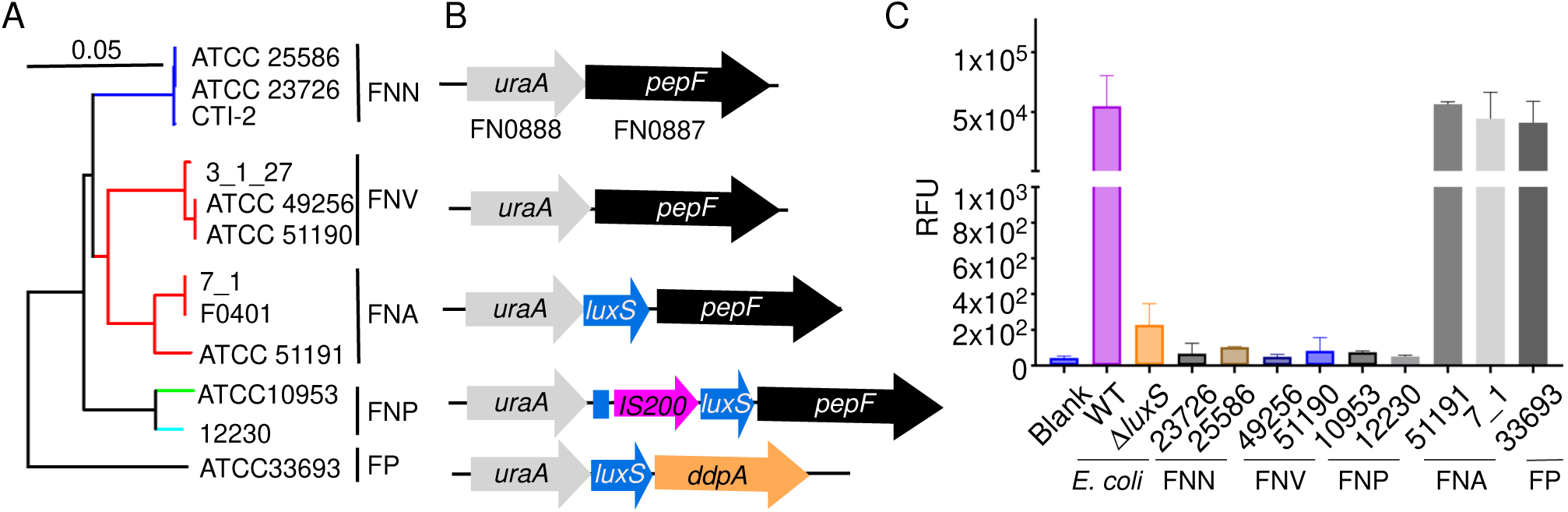
AI-2 production is restricted to *F. nucleatum* subsp. *animalis* and correlates with an intact *luxS* locus. (A) Phylogenetic relationship of representative *F. nucleatum* strains used in this study, grouped by subspecies—subsp. *nucleatum* (FNN; ATCC 25586, ATCC 23726, CTI-2), subsp. *vincentii* (FNV; 3_1_27, ATCC 49256, ATCC 51190), subsp. *animalis* (FNA; 7_1, F0401, ATCC 51191), and subsp. *polymorphum* (FNP; ATCC 10953, 12230). The phylogenetic tree was constructed based on znpA gene using the maximum-likelihood method implemented in DNAMAN Version 10 (Lynnon Biosoft). *Fusobacterium periodonticum* ATCC 33693 (FP) was included as an outgroup. (B) Schematic of the chromosomal region between *uraA* and *pepF* showing subspecies-specific presence of *luxS*. *luxS* is absent from FNN and FNV at this locus, present as an intact gene in FNA (between *uraA* and *pepF*), and disrupted in FNP by insertion of an IS200-family element. The corresponding region from *F. periodonticum* is shown for comparison. Arrows indicate gene orientation; *uraA* (gray), *pepF* (black), *luxS* (blue), IS200 insertion (magenta), and the adjacent gene (*ddpA*, orange) are indicated. (C) AI-2 activity in cell-free culture supernatants was measured using the *Vibrio harveyi* BB170 bioluminescence reporter assay. Supernatants from FNN, FNV, and FNP strains showed signals at or near background levels, whereas all tested FNA strains and *F. periodonticum* generated robust reporter induction. *E. coli* wild type (WT) and its Δ*luxS* mutant served as positive and negative controls, respectively. Data are presented as relative fluorescence units (RFU; mean ± SD) from three independent experiments (each assayed in technical triplicate); the y-axis includes a break to display both low- and high-signal samples.

To test this prediction experimentally and reassess prior reports, we measured AI-2 production in representative strains from each subspecies using the *Vibrio harveyi* BB170 bioluminescence assay (see Material and Method). Consistent with the genomic analysis, FNN, FNV, and FNP strains did not produce detectable AI-2 under the conditions tested (Fig. 1C). In contrast, FNA strains produced robust AI-2 signals comparable to those observed in the positive control *Escherichia coli* and in *Fusobacterium periodonticum*, a closely related species that we previously showed to produce AI-2(41).

To directly determine whether the absence of AI-2 in FNN strains is solely due to the lack of *luxS*, we used FNN strain ATCC 23726, a genetically tractable and well-established model strain for *F. nucleatum* research (42). Because this strain can be readily engineered, it allowed us to test whether the introduction of *luxS* is sufficient to confer AI-2 production. We inserted the *luxS* gene from FNA strain 7_1, together with its native ribosome-binding site, into the chromosome between *uraA* and *pepF* (Fig.2A & 2B). The engineered strain produced AI-2 at levels comparable to FNA 7_1 and *E. coli* (Fig. 1C & 2C), demonstrating that FNN strains retain the metabolic capacity to produce AI-2 when supplied with a functional *luxS* gene. The FNN strain ATCC 25586, previously reported to produce AI-2, is difficult to genetically manipulate(43). To evaluate this strain, we introduced a shuttle plasmid that expresses FNA 7_1 luxS under the control of the *pfdx* promoter(44). AI-2 production was detected only in the strain carrying the *luxS*-expressing plasmid and not in the parental strain (Fig.2C). Notably, expression of *luxS* and restoration of AI-2 production did not alter bacterial growth or monospecies biofilm formation (Fig. 2D & 2E & Fig.S1). Together, these results demonstrate that AI-2 production in *F. nucleatum* depends on an intact *luxS* gene and is restricted to FNA. AI-2 production is therefore a subspecies-specific metabolic trait rather than a conserved property of the species.

**Figure 2.**
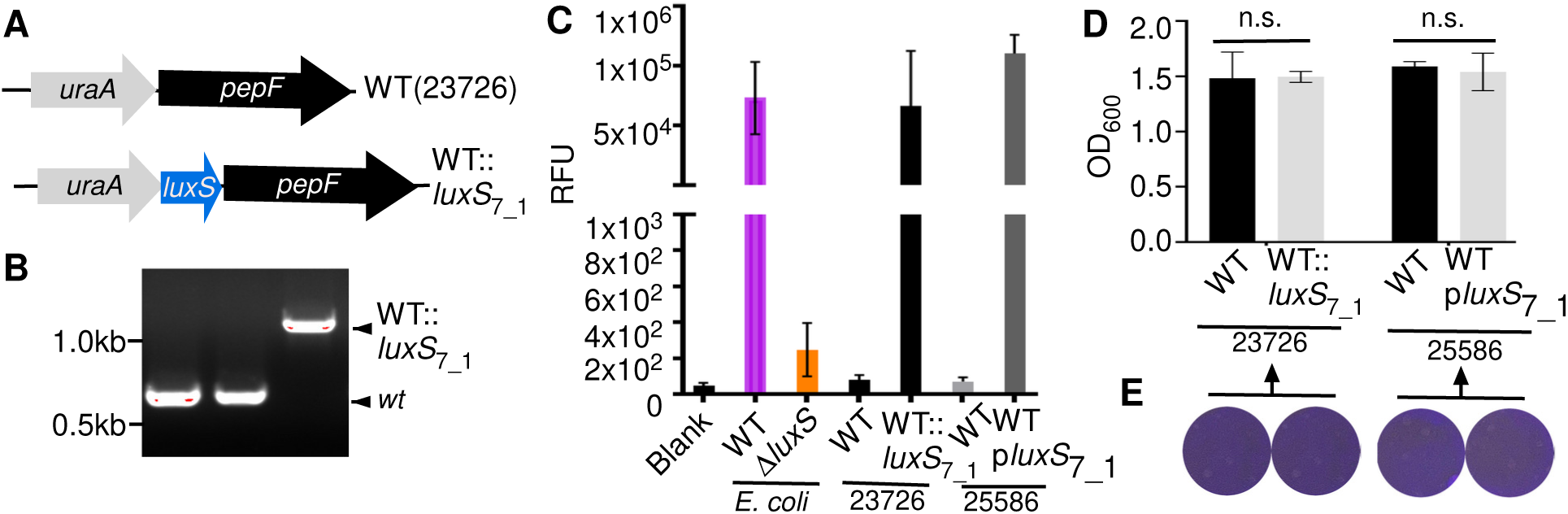
Introduction of *luxS* from FNA strain 7_1 restores AI-2 production in FNN strains without detectable effects on growth or monospecies biofilm formation. **(A)** Schematic of the *uraA–pepF* chromosomal locus in FNN ATCC 23726 before (WT) and after in-frame insertion of the FNA 7_1 *luxS* gene (WT::*luxS*_7_1_) between *uraA* and *pepF*. **(B)** PCR confirming correct chromosomal insertion of *luxS*7_1 in ATCC 23726 (WT band versus the larger amplicon from WT::*luxS*7_1). **(C)** AI-2 activity measured by the *V. harveyi* BB170 bioluminescence reporter assay. WT ATCC 23726 and WT ATCC 25586 showed background-level signals, whereas ATCC 23726 carrying the chromosomal *luxS*_7_1_ insertion (WT::*luxS*_7_1_) and ATCC 25586 expressing *luxS*_7_1_ from a shuttle plasmid (p*luxS*_7_1_) produced robust AI-2 signals comparable to the *E. coli* WT positive control. The *E. coli* Δ*luxS* strain served as a negative control. **(D)** Growth analysis showing that expression of *luxS*7_1 in ATCC 23726 (WT::*luxS*_7_1_) or ATCC 25586 (p*luxS*_7_1_)) did not significantly alter final culture density relative to the corresponding WT strains (n.s., Student’s *t* test). **(E)** Representative crystal violet–stained monospecies biofilms demonstrating no obvious difference in biofilm biomass between WT and *luxS*_7_1_-expressing derivatives of ATCC 23726 and ATCC 25586 after anaerobic growth in TSPC for 72 h.

### *luxS* Is Dispensable for Growth and Biofilm Formation in FNA

Because AI-2 production is restricted to FNA, we next asked whether LuxS-dependent AI-2 synthesis provides a physiological advantage to this subspecies. If AI-2 functions as a quorum-sensing regulator, its loss would be expected to affect growth, fitness, or biofilm development (1, 45). To test this directly, we deleted *luxS* in an FNA strain and examined the resulting phenotype.

We chose FNA strain 7_1, a well-characterized human gut isolate that is widely used as a representative FNA strain (46–48). An in-frame deletion of *luxS* was generated using our previously developed conditional plasmid system for gene deletion in genetically recalcitrant *F. nucleatum* strains(41). The deletion plasmid (pBCG10Δ*luxS*) carries flanking regions upstream and downstream of *luxS* to facilitate homologous recombination (Fig. 3A). For counterselection, we used the toxin gene *mazF* (49), which eliminates cells retaining the plasmid during allelic exchange (see Materials and Methods). Using this strategy, we successfully obtained a Δ*luxS* mutant (Fig.3B). AI-2 production was completely abolished in the mutant, as measured by the *V. harveyi* BB170 bioluminescence assay (Fig. 3C), confirming that *luxS* is solely responsible for AI-2 synthesis in FNA. Expression of *luxS* in trans from a plasmid (p*LuxS*) restored AI-2 production to wild-type levels, validating the specificity of the deletion (Fig.3C).

**Figure 3.**
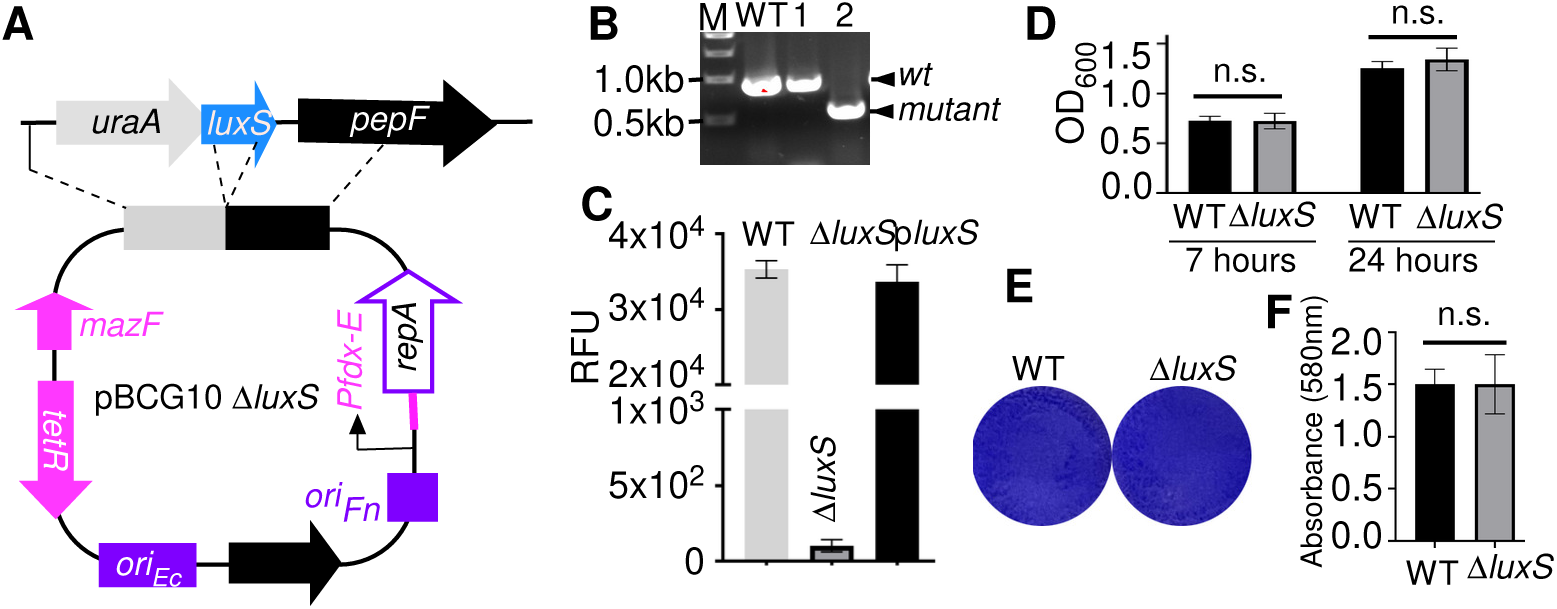
Deletion of *luxS* in FNA strain 7_1 abolishes AI-2 production but does not affect growth or monospecies biofilm formation. **(A)** Schematic representation of the conditional suicide plasmid pBCG10-Δ*luxS* used for in-frame deletion of *luxS*. Plasmid replication in *F. nucleatum* is controlled by a theophylline-responsive riboswitch (Pfdx-E) regulating *repA*. The toxin gene *mazF* is under the control of the ATc-inducible TetR system for counterselection. Origins of replication for *E. coli* (oriEC) and *F. nucleatum* (oriFN) are indicated. Flanking regions upstream and downstream of *luxS* enable homologous recombination. **(B)** Diagnostic PCR confirming successful generation of the Δ*luxS* mutant in strain 7_1. **(C)** AI-2 activity measured using the *V. harveyi* BB170 reporter assay. Deletion of *luxS* completely abolished AI-2 production, whereas complementation in trans restored AI-2 activity to wild-type levels. Data are presented as relative fluorescence units (RFU; mean ± SD) from three independent experiments, each performed in technical triplicate. **(D)** Growth analysis showing no significant difference between WT and Δ*luxS* strains at the indicated time points (n.s., Student’s *t* test). **(E)** Representative crystal violet–stained monospecies biofilms after 3 days of anaerobic growth in TSPC medium, demonstrating no apparent difference between WT and Δ*luxS* strains. **(F)** Quantification of biofilm biomass measured by crystal violet staining. Values represent the mean ± SD from three independent experiments performed in triplicate. Statistical analysis was conducted using Student’s *t-test* in GraphPad Prism.

We then examined whether loss of *luxS* affects bacterial growth. Under standard anaerobic conditions, the Δ*luxS* mutant displayed growth indistinguishable from the wild type (Fig. 3D). Thus, LuxS is not required for normal growth in FNA. Because AI-2 has previously been implicated in biofilm formation in *F. nucleatum*, we next assessed monospecies biofilm development. Wild-type and Δ*luxS* strains were grown anaerobically for 72 hours in 12-well plates containing TSPC medium adjusted to pH 8.5, a condition known to promote biofilm formation(50, 51). Biofilms were quantified by crystal violet staining(52). No significant difference in biofilm biomass was observed between the wild type and the Δ*luxS* mutant (Fig. 3E & 3F). Taken together, these results demonstrate that although AI-2 production is a defining metabolic feature of FNA, LuxS-dependent AI-2 synthesis is dispensable for both growth and monospecies biofilm formation under the conditions tested.

### Deletion of *luxS* Does Not Induce a Quorum-Sensing–Like Transcriptional Response

Because deletion of *luxS* did not affect growth or monospecies biofilm formation in FNA, we next asked whether AI-2 regulates gene expression at the transcriptional level, as expected for a canonical quorum-sensing system. In classical AI-2–mediated signaling, accumulation of the molecule at high cell density typically triggers substantial transcriptional reprogramming, often involving large (more than 10–100 fold) changes in target gene expression(45, 53, 54). If AI-2 functions as a quorum-sensing signal in FNA, deletion of *luxS* should therefore produce broad and robust transcriptional alterations.

To examine this, we first monitored extracellular AI-2 levels during growth of FNA strain 7_1. AI-2 accumulated during exponential growth, reached a peak in late log phase, and then decreased slowly thereafter. As expected, the FNN strain ATCC 23726, which lacks *luxS*, did not produce detectable AI-2 at any growth stage (Fig.4A). Guided by these kinetics, we collected RNA from wild-type 7_1 and the Δ*luxS* mutant at two physiologically relevant growth stages—one before peak AI-2 accumulation and one after. Differential expression analysis was performed using a ≥2-fold cutoff (|log2 fold change| ≥ 1).

**Figure 4.**
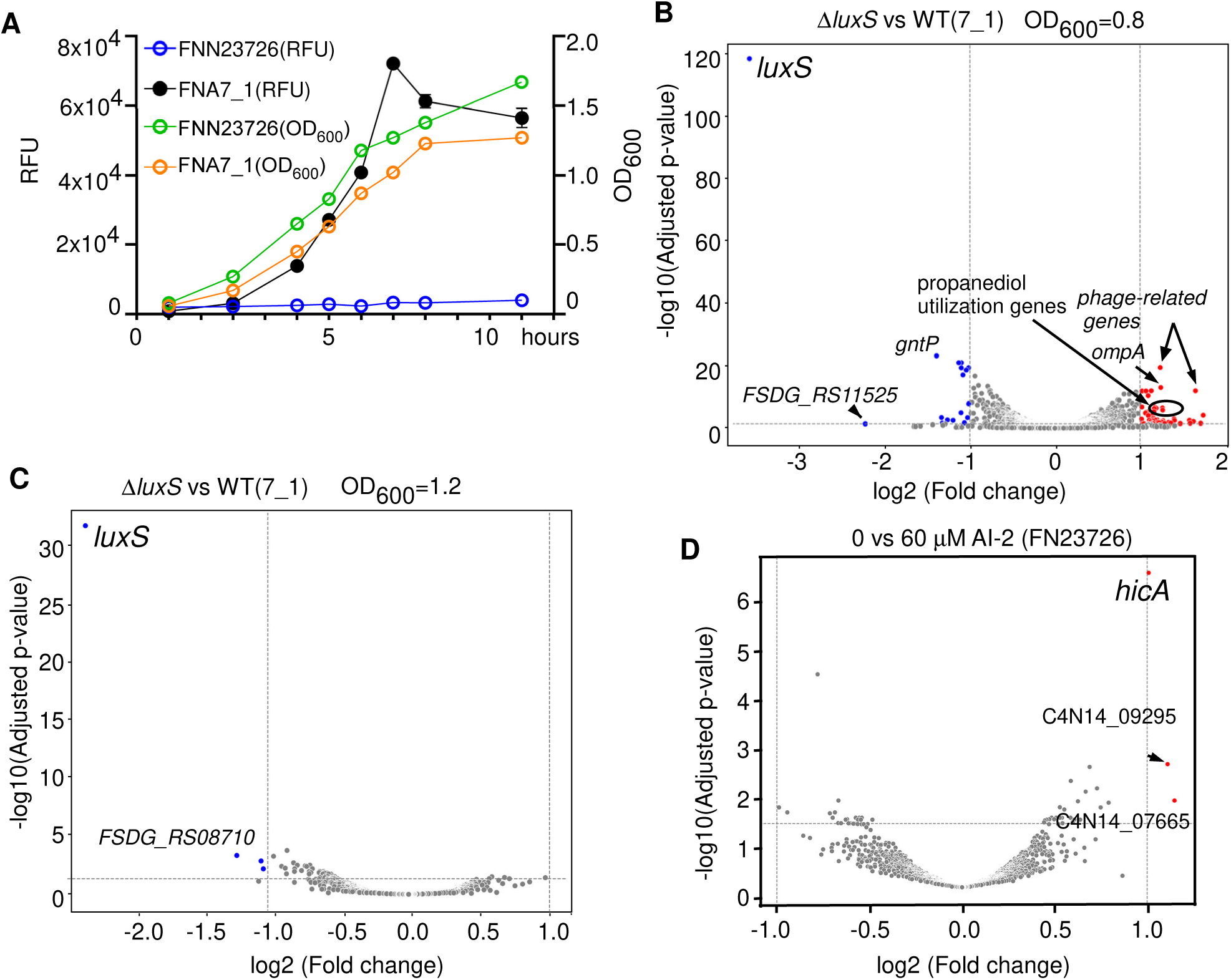
Deletion of *luxS* in FNA strain 7_1 results in minimal transcriptional changes. **(A)** Time-course analysis of extracellular AI-2 activity during growth of FNA strain 7_1. AI-2 levels were measured using the *V. harveyi* BB170 reporter assay, and culture density (OD_600_) was recorded at each time point. FNN ATCC 23726, which lacks *luxS*, served as a negative control. Data represent the mean ± SD from three independent experiments. **(B)** Volcano plot showing differential gene expression between Δ*luxS* and WT 7_1 at OD_600_ ≈ 0.8, as determined by RNA-seq analysis. Genes meeting the threshold of |log₂(fold change)| ≥ 1 and *p* ≤ 0.05 are highlighted. **(C)** Volcano plot showing differential gene expression between Δ*luxS* and WT 7_1 at OD_600_ ≈ 1.2. Few genes met the differential expression threshold, and fold changes were modest in magnitude. **(D)** Volcano plot of RNA-seq analysis comparing FNN ATCC 23726 treated with synthetic AI-2 (DPD) versus untreated controls. Only a small number of genes showed ≥2-fold changes, indicating that exogenous AI-2 does not induce a coordinated transcriptional response in this subspecies.

At the earlier point, deletion of *luxS* resulted in 71 genes showing increased expression and 23 genes showing decreased expression relative to the wild type. However, aside from *luxS* itself, which showed the expected strong reduction, the magnitude of differential expression was modest, generally limited to 2- to 3-fold changes (Fig. 4B and Table S3). Among the upregulated genes were two phage-associated genes and two genes related to propanediol utilization, but no coherent regulatory pattern consistent with a quorum-sensing regulon was observed. Importantly, these changes were not sustained. At the later growth stage, no genes were significantly upregulated in the Δ*luxS* mutant, and only five genes—including *luxS*—were downregulated, again with small fold changes (Fig. 4C & Table S4).

We next tested whether non-AI-2–producing subspecies might still sense and respond to extracellular AI-2. FNN strain ATCC 23726 was treated with 30 μM synthetic DPD, a concentration that induces *V. harveyi* BB170 reporter activity comparable to the peak AI-2 levels detected in late-log–phase FNA culture supernatants (Fig. S2). After 1.5 hours of exposure, RNA-seq analysis revealed only three genes with ≥2-fold increased expression and no significantly downregulated genes. The observed changes were minor and lacked a coordinated regulatory pattern (Fig. 4D & Table S5). Together, these transcriptomic results indicate that AI-2 does not drive global gene expression changes in *F. nucleatum*, suggesting that LuxS-dependent AI-2 production is not coupled to a functional quorum-sensing regulatory system.

### The Activated Methyl Cycle Is Essential for Viability in *F. nucleatum*

LuxS functions within the activated methyl cycle (AMC), where it participates in recycling S-adenosylhomocysteine (SAH) generated during methylation reactions (Fig. 5A). Although deletion of *luxS* abolished AI-2 production, it did not impair growth or biofilm formation in FNA, and the other three subspecies either lack *luxS* entirely or carry a disrupted allele. In many bacteria that do not encode LuxS, the AMC is maintained through an alternative pathway mediated by S-adenosylhomocysteine hydrolase (SahH, also known as AhcY), which directly converts SAH into homocysteine and bypasses the LuxS-dependent step(55, 56). However, comprehensive genome searches across all four *F. nucleatum* subspecies failed to identify any *sahH* homolog. This absence raised a fundamental question: is the activated methyl cycle itself essential for viability in *F. nucleatum*, and how is it maintained in subspecies lacking functional LuxS and SahH?

**Figure 5.**
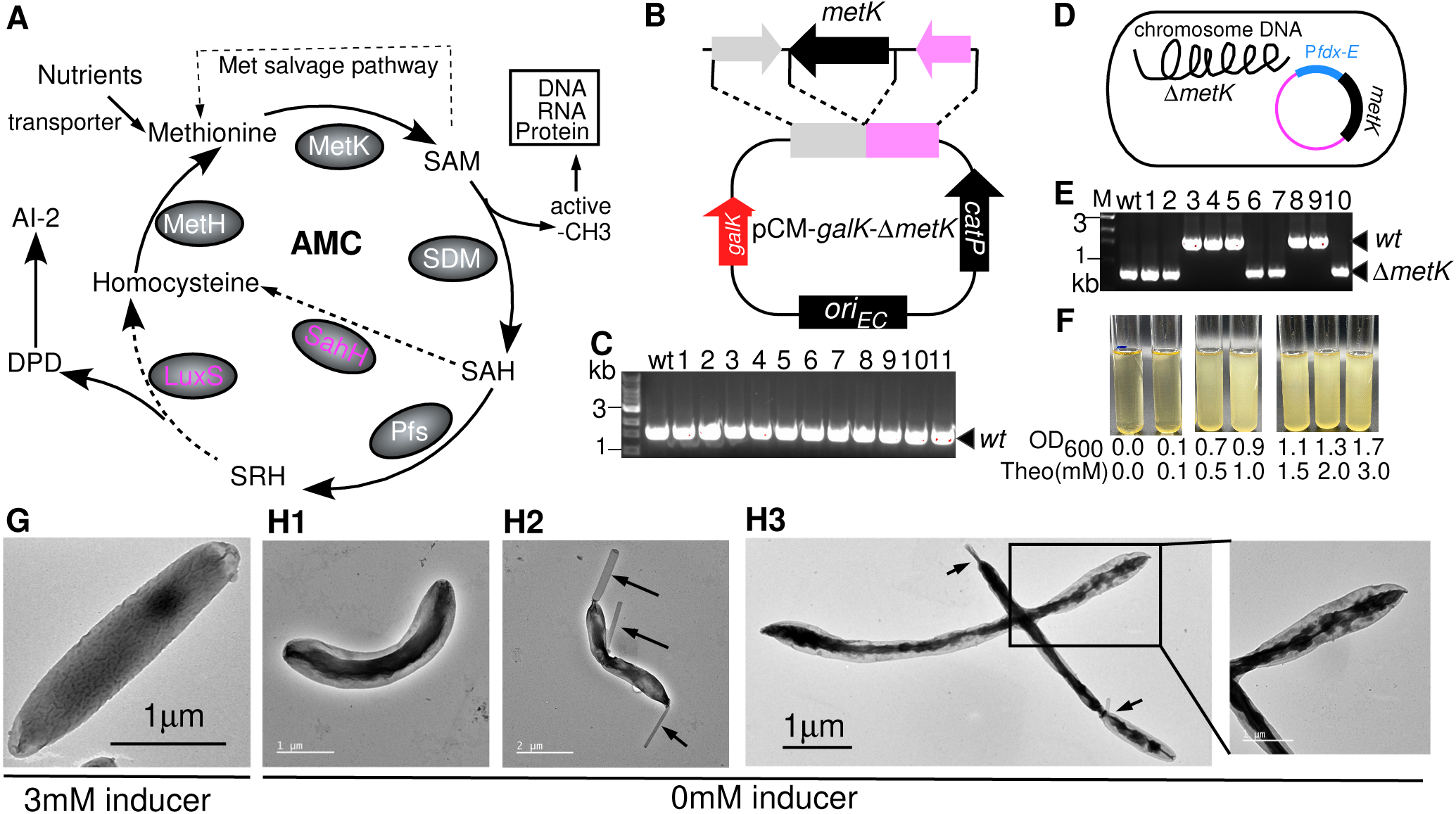
The activated methyl cycle (AMC) is essential for viability in *Fusobacterium nucleatum*. **(A)** Schematic representation of the activated methyl cycle (AMC) illustrating methyl group transfer and methionine recycling. MetK (S-adenosylmethionine synthetase) converts methionine to S-adenosylmethionine (SAM), the universal methyl donor. Following methyl transfer by S-adenosylmethionine-dependent methyltransferases (SDMs), SAM is converted to S-adenosylhomocysteine (SAH). In the LuxS-dependent pathway, SAH is processed by Pfs (5′-methylthioadenosine/S-adenosylhomocysteine nucleosidase) to form S-ribosylhomocysteine (SRH), which is subsequently cleaved by LuxS to generate homocysteine and the AI-2 precursor DPD. In alternative pathways found in other bacteria, SAH can be directly converted to homocysteine by SahH (S-adenosylhomocysteine hydrolase). **(B)** Schematic diagram of the *metK* deletion construct (pCM-galK-Δ*metK*) used to generate an in-frame chromosomal deletion in FNN ATCC 23726. Approximately 1.5 kb of upstream and downstream homologous regions flank the deleted *metK* coding sequence to facilitate double-crossover recombination. (**C)** PCR screening of more than 100 counterselected colonies following allelic exchange showed retention of the wild-type *metK* allele, with no Δ*metK* mutants recovered, indicating that *metK* is essential under the tested conditions. Representative PCR results from 10 independent colonies are shown. **(D)** Strategy for the construction of a conditional *metK* mutant. Because *metK* is essential, chromosomal deletion was performed in the presence of a plasmid-borne copy of *metK* expressed under the control of a theophylline- inducible riboswitch, allowing complementation in trans. **(E)** PCR confirmation of successful chromosomal deletion of *metK* in the presence of plasmid-mediated complementation, demonstrating that deletion is possible only when *metK* expression is provided in trans. **(F)** Growth analysis of the conditional *metK* mutant showing strict dependence on theophylline for viability. Bacterial growth exhibited a dose-dependent response to the inducer, and no growth was observed in its absence, confirming that *metK* is essential for survival in *F. nucleatum*. **(G)** Transmission electron microscopy (TEM) of the conditional Δ*metK* strain. Cells grown in the presence of 3 mM theophylline displayed normal morphology comparable to wild type. In contrast, depletion of *metK* (no inducer; cells precultured with 2 mM theophylline and then grown for 12 h without inducer) resulted in pronounced morphological abnormalities, including curved cells (**H1**), surface-associated tubular-like structures (**H2**), and marked cell elongation (H3; enlarged view shown).

The AMC is initiated by MetK, the S-adenosylmethionine synthetase that converts methionine into S-adenosylmethionine (SAM), the universal methyl donor required for DNA, RNA, protein, and metabolite methylation (Fig. 5A). In *E. coli* and many other bacteria, *metK* is essential because methyl metabolism is indispensable for cellular function(57, 58). To determine whether AMC activity is essential in *F. nucleatum,* we attempted to delete *metK* in the genetically tractable FNN strain ATCC 23726.

A deletion construct (pCM-*galK*-Δ*metK*) containing flanking regions of *metK* was introduced into ATCC 23726, and plasmid integration was obtained (Fig. 5B). To resolve the integrated plasmid, cultures were grown without antibiotic selection and subsequently plated on TSPC agar containing 0.25% 2-deoxy-D-galactose (2-DG) to select for second crossover events. In this galK-based counterselection system, reconstitution of *galK* renders cells sensitive to 2-DG; thus, 2-DG-resistant colonies represent candidates that have lost the plasmid backbone following recombination. If *metK* were nonessential, we would expect recovery of both wild-type revertants and Δ*metK* mutants at approximately equal frequency. However, after screening more than 100 counterselected colonies, no Δ*metK* mutants were recovered; all isolates retained the wild-type allele (Fig. 5C). These results strongly suggest that *metK* is essential for viability in *F. nucleatum*.

To directly confirm this, we introduced an inducible copy of *metK* on a shuttle plasmid under control of a theophylline-responsive riboswitch. In the presence of the inducer, chromosomal deletion of *metK* was successfully achieved, demonstrating that viability can be supported by plasmid-borne expression (Fig. 5D & 5E). Growth of the resulting strain was strictly dependent on theophylline, with a dose-dependent effect. In the absence of the inducer, cells failed to grow (Fig. 5F). At 3 mM theophylline, cell morphology was indistinguishable from wild type (Fig.5G). In contrast, upon *metK* depletion, cells exhibited pronounced morphological abnormalities, including curvature (Fig. 5H1), the formation of tubular-like structures (Fig. 5H2 and 5H3, arrows), elongation, and severe membrane damage, ultimately leading to cell death (Fig. 5H3). Together, these results show that while AI-2 production is dispensable and not linked to quorum sensing, the activated methyl cycle is essential for *F. nucleatum* viability, indicating that AI-2 in FNA is a retained metabolic byproduct rather than a regulatory signal.

## DISCUSSION

In this study, we demonstrate that AI-2 production in *F. nucleatum* is subspecies-specific and restricted to subsp. *animalis*, where an intact *luxS* gene is retained. The other three subspecies either lack *luxS* entirely (FNN and FNV) or harbor a disrupted allele (FNP). Functional analyses showed that deletion of *luxS* in FNA abolishes AI-2 production but does not affect growth, biofilm formation, or global gene expression. Moreover, exogenous AI-2 fails to induce a transcriptional response in non-producing subspecies. Finally, we demonstrate that while LuxS-dependent AI-2 production is dispensable, the upstream activated methyl cycle (AMC) is essential for viability, as *metK* cannot be deleted without conditional complementation. Together, these findings establish that AI-2 production in *F. nucleatum* is not coupled to a canonical quorum-sensing regulatory system.

Our findings differ from earlier reports, suggesting that FNN and FNP strains, including ATCC 25586 and ATCC 10953, produce AI-2 and utilize it in monospecies and polymicrobial biofilm formation(7, 34–39). In addition, a series of studies from a same group reported AI-2 production in *F. nucleatum* strain F01, which, based on 16S rRNA analysis, appears to belong to FNP; however, because its genome sequence is not available, the integrity of its *luxS* locus cannot be evaluated(59–63). Importantly, we employed the same *V. harveyi* BB170 reporter assay used in previous studies but observed no detectable AI-2 activity in FNN or FNP strains when measuring directly from filtered culture supernatants. In contrast, some prior reports relied on partially purified preparations involving size-exclusion filtration and C18 chromatography before assay(34, 59). The enrichment requirement suggests that the detected signals were low in abundance and may have been influenced by other small molecules present in the complex media.

It is well established that the BB170 assay can be influenced by residual nutrients and trace components(64). For example, elevated glucose concentrations can enhance reporter growth and amplify luminescence through endogenous AI-2 self-induction, and trace elements such as borate can affect signal intensity(65). Differences in culture media composition may therefore contribute to artifactual or background-level signals that are difficult to distinguish from true LuxS-dependent AI-2 production. Importantly, even partially purified preparations may retain low–molecular–weight nutrients or metabolic intermediates that stimulate BB170 growth and artificially enhance luminescence, potentially generating false-positive signals. In our experiments, the signal levels previously reported for ATCC 10953 were comparable to background readings(7). Taken together, differences in assay conditions, media composition, and sample processing provide plausible explanations for the discrepancy between our results and earlier studies. When combined with our genomic analyses demonstrating the absence or disruption of *luxS* in FNN and FNP, our data strongly support the conclusion that AI-2 production is restricted to subsp. *animalis*.

FNN, FNV, and FNP strains do not produce detectable AI-2 (Fig.1), and exogenous AI-2 addition does not alter gene expression in FNN (Fig.3D). Although FNA produces AI-2, deletion of luxS abolishes AI-2 synthesis yet causes only minimal transcriptional changes (Fig.3B & 3C). Together, these findings indicate that *F. nucleatum* neither senses nor utilizes AI-2 as a canonical quorum-sensing signal(66). Consistent with this conclusion, we did not identify homologs of known AI-2 receptor proteins, such as LsrB in *E. coli* or LuxP in *Vibrio* species(56, 67, 68), in any *F. nucleatum* genome.

The absence of AI-2–mediated quorum sensing may reflect the ecological lifestyle of *F. nucleatum*. Unlike bacteria that alternate between planktonic growth and biofilm formation, *F. nucleatum* is rarely solitary in the oral cavity. Instead, it rapidly integrates into polymicrobial biofilms, where it serves as a structural bridge linking early and late colonizers(69, 70). In this environment, local cell density is persistently high. Classical quorum-sensing systems function as density-dependent switches, activating gene expression only when population density reaches a threshold. However, if *F. nucleatum* is constitutively present in high-density biofilms, an AI-2–mediated regulatory circuit would be chronically activated, thereby losing its adaptive value as a conditional response mechanism.

Importantly, the lack of AI-2 production in non-FNA strains does not diminish its ecological importance. Non-FNA strains, especially FNN, are the dominant architectural components of dental plaque due to their exceptionally strong coaggregation capacity(70–72). Many neighboring oral bacteria, including streptococci and *P. gingivalis*, produce AI-2(69). By physically bringing these organisms into close proximity, non-FNA facilitates interspecies interactions—including AI-2–mediated communication among other species—without producing the signal itself. Its structural role alone is sufficient to promote community-level signaling within the oral biofilm.

This raises the question of why FNA retains an intact *luxS* gene. Compared with other subspecies, FNA generally exhibits weaker coaggregation with oral bacteria, such as Actinomyces oris (Fig. S3), and may exhibit more dynamic spatial organization within host tissues. Rather than mediating intraspecies quorum sensing, AI-2 production in FNA may influence host-associated processes. If LuxS-dependent AI-2 production enhances virulence or host adaptation, this hypothesis should be directly tested in future animal infection models. The selective retention of *luxS* in FNA may therefore reflect subspecies-specific adaptation rather than quorum-sensing regulation.

Another important question raised by our findings concerns the functionality of the activated methyl cycle (AMC) in *F. nucleatum*. FNN, FNV, and FNP lack an intact *luxS* gene, and deletion of *luxS* in FNA does not impair growth. This raises the question of how methyl group recycling and methionine regeneration are maintained in this species. In many bacteria, S-adenosylhomocysteine hydrolase (SahH, also known as AhcY) provides an alternative route for recycling S-adenosylhomocysteine (SAH), thereby bypassing the LuxS-dependent step of the AMC(55). However, genome-wide searches across all *F. nucleatum* subspecies failed to identify a *sahH* homolog. The combined absence of SahH and the variable retention of *luxS* suggest that AMC flux and methionine metabolism in *F. nucleatum* may be adapted to its specific ecological niche.

Despite the apparent incompleteness of the canonical AMC, we demonstrated that *metK*, which encodes S-adenosylmethionine synthetase, is essential for viability (Fig. 5). This indicates that methyl metabolism itself is indispensable. However, in the absence of both LuxS (in most subspecies) and SahH, the classical AMC pathway would not efficiently regenerate methionine or recycle methyl groups internally. One possible explanation is that *F. nucleatum* relies heavily on environmental methionine uptake to satisfy its metabolic demands. Previous studies have shown that *F. nucleatum* can import methionine(73, 74). Methionine could be obtained either through the methionine salvage pathway(75, 76) or directly from the environment. In the subgingival pocket, both *F. nucleatum* and its co-colonizer *P. gingivalis* secrete proteases that degrade host tissues, releasing amino acids, including methionine. Continuous access to exogenous methionine may therefore compensate for incomplete internal recycling of the AMC. If this model is correct, accumulation of S-ribosylhomocysteine (SRH), the LuxS substrate, may occur in strains lacking functional *luxS*. Expression of *luxS* in non-FNA strains would be expected to reduce extracellular SRH levels. We are currently quantifying SRH concentrations in wild-type and *luxS*-expressing strains using mass spectrometry. If no difference in SRH levels is observed, this would suggest the existence of an alternative, as yet unidentified, enzyme participating in methyl-cycle processing in *F. nucleatum*.

In FNA, the retention of *luxS* may reflect adaptation to distinct ecological pressures. If FNA preferentially colonizes environments where methionine availability is limited, maintenance of a functional LuxS-dependent branch of the activated methyl cycle (AMC) could confer a selective advantage by facilitating SAH detoxification and methyl flux balance. Notably, free methionine levels in the human intestinal lumen—where FNA is frequently detected in diseased states—are reported to be very low(77, 78). Therefore, variation in methionine availability across host niches may help explain the subspecies-specific retention of *luxS*.

Taken together, our findings challenge the prevailing view that AI-2 functions as a universal quorum-sensing signal in *F. nucleatum*. Rather than serving as a density-dependent regulatory system, AI-2 production in this species appears to reflect subspecies-specific metabolic adaptation. The differential retention of *luxS* among subspecies highlights the importance of ecological context in shaping metabolic pathways and cautions against assuming conserved signaling mechanisms across genetically diverse lineages. More broadly, this work highlights the importance of examining subspecies-level variation when interpreting signaling phenotypes in complex microbial communities.

## MATERIALS AND METHODS

All bacterial strains and plasmids employed in this work are summarized in Table S1. *F. nucleatum* and *F. periodonticum* strains were propagated either in liquid medium or on solid medium consisting of tryptic soy broth supplemented with 1% (w/v) Bacto Peptone and 1 mM freshly prepared cysteine (designated TSPC). For plate cultivation, the same formulation was solidified with agar. Anaerobic growth was carried out in a Coy anaerobic chamber (Coy Laboratory Products) containing a gas atmosphere of N₂, H₂, and CO₂. The hydrogen concentration within the chamber was routinely monitored and maintained above 2% using a portable H₂ detector positioned inside the workstation. *E. coli* strains were cultured aerobically in Luria–Bertani (LB) medium or on LB agar plates. Where appropriate, antibiotics were supplemented at the following final concentrations: chloramphenicol (15 μg/mL), thiamphenicol (5 μg/mL), erythromycin (300 μg/mL in LB agar and 50 μg/mL in LB broth), and clindamycin (1 μg/mL).

### Plasmids construction

The oligonucleotide primers used in this study were synthesized by Sigma-Aldrich and are listed in Table S2. Plasmid constructs generated in this work were assembled using the Gibson assembly method with the NEBuilder HiFi DNA Assembly Master Mix (New England Biolabs, E2621L), according to the manufacturer’s instructions. All recombinant plasmids were confirmed by DNA sequencing to ensure sequence accuracy.

i. pBCG02-*luxS_7_1_* — This plasmid was constructed to integrate the *luxS* gene from FNA strain 7_1 into the chromosome between *uraA* and *pepF* of FNN ATCC 23726. To generate the upstream homologous arm, a 1.5-kb fragment immediately upstream of the *uraA* stop codon in ATCC 23726 was amplified using primers uraA-upF and uraA-upR. The downstream homologous arm consisted of a 1.5-kb region located downstream of the *uraA* stop codon, amplified with primers pepF-dnF and pepF-dnR. The *luxS* gene, including its native ribosome-binding site and spanning from the start to the stop codon, was amplified from strain 7_1 and used as the middle fragment. The upstream, middle (*luxS*), and downstream fragments were assembled into the suicide vector pBCG02(79), which had been linearized by PCR using primers BCG02Mch-F and BCG02Mch-R. Gibson assembly was employed to complete the construction.
ii. p*Lux*_7_1_ —This plasmid was generated to express the *luxS* gene from *FNA* strain 7_1 in trans under the control of the *uraA* promoter using the shuttle vector pBCG11(80). The promoter region of *pfdx* was amplified from pCWU50 with primers pfdx-F and pfdx-R. The *luxS* coding sequence, including its native ribosome-binding site and approximately 100 bp downstream of the stop codon, was amplified from strain 7_1 using primers luxS7_1F2 and luxS7_1R2. The pBCG11 backbone was PCR-linearized with primers pBCG11-F and pBCG11-R. The *uraA* promoter fragment and the *luxS* insert were then assembled into the linearized vector by Gibson assembly, generating plasmid pLux7_1.
iii. pBCG10-*ΔluxS* — The conditional suicide plasmid pBCG10 was first generated by modifying pZP06C(41). Briefly, the *sacB* counterselection cassette driven by the *PrpsJ* promoter was removed by PCR amplification of the pZP06C backbone excluding the *PrpsJ-sacB* region using primers pZP06-nosacB-F and pZP06-nosacB-R. The *tetR-mazF* cassette was amplified from plasmid pVoPo-04(49) using primers tetR-mazF-F and tetR-mazF-R, which introduced overlapping sequences compatible with the linearized backbone. Gibson assembly was performed to replace *sacB* with the ATc-inducible *mazF* toxin module, yielding pBCG10. In this construct, plasmid replication in *Fusobacterium* is controlled by a theophylline-responsive riboswitch regulating *repA*, whereas *mazF* expression is governed by the tetracycline-inducible TetR system. To construct pBCG10-Δ*luxS* for the deletion of *luxS* in FNA strain 7_1, approximately 1.5-kb regions upstream and downstream of *luxS* were amplified from genomic DNA using primer pairs luxSupF/R and luxSdnF/R, respectively. The native *luxS* gene encodes a 159-amino-acid protein; in this design, an internal in-frame deletion spanning amino acids 10–121 was introduced, removing the central catalytic region while preserving the flanking sequences. The pBCG10 backbone was PCR-linearized using primers BCG10F and BCG10R, and the upstream and downstream homologous arms were assembled into the linearized vector by Gibson assembly. The resulting construct, pBCG10-Δ*luxS*, contains the fused flanking regions and is designed to mediate double-crossover homologous recombination to generate an in-frame Δ*luxS* mutant.
iv. pCM-galK-Δ*metK* — To construct the *metK* deletion plasmid, approximately 1.5-kb regions upstream and downstream of *metK* were amplified from the genomic DNA of FNN ATCC 23726 using primer pairs metKupF/metKupR and metKdnF/metKdnR, respectively. The upstream fragment was digested with *SacI* and *KpnI*, whereas the downstream fragment was digested with *KpnI* and *SalI*. These fragments were sequentially ligated into the vector pCM-*galK* that had been pre-digested with *SacI* and *SalI*, generating plasmid pCM-galK-Δ*metK*. The integrity of the resulting construct was verified by colony PCR and restriction enzyme analysis.
v. pSX11E-P*fdx-E-metK* — This plasmid was constructed to express a functional copy of *metK* in trans from an extrachromosomal location, thereby permitting subsequent deletion of the chromosomal *metK* gene. The shuttle vector pBCG11 was initially available; however, both pBCG11 and the deletion plasmid pCM-galK carry the *catP* gene conferring thiamphenicol resistance, precluding their simultaneous use. To resolve this incompatibility, a modified pBCG11-derived vector was generated in which the *catP* cassette was replaced with the *ermF/AM* resistance determinant, conferring clindamycin resistance. To construct this intermediate plasmid (designated pSX11E), the pBCG11 backbone lacking *catP* was amplified using primers re-pBCG11F and re-pBCG11R. The *ermF/AM* cassette was amplified from plasmid pG106 using primers ermFAM-F and ermFAM-R. The linearized backbone and the resistance cassette were assembled by Gibson assembly, generating pSX11E. For controlled expression of *metK*, the gene was placed under the control of the *Pfdx-E* regulatory module, which consists of the constitutive *fdx* promoter coupled to a theophylline-responsive riboswitch previously validated for inducible gene expression in *Fusobacterium nucleatum*. The pSX11E vector was PCR-linearized, and the *Pfdx-E* fragment was amplified from plasmid pZP4C using primers pfdx-metKF and pfdx-metKR. The *metK* open reading frame was amplified from *F. nucleatum* subsp. *nucleatum* ATCC 23726 genomic DNA using primers com-metKF and com-metKR. The linearized vector, *Pfdx-E* fragment, and *metK* coding sequence were assembled by Gibson assembly, yielding plasmid pSX11E-Pfdx-E-metK.

### In-frame insertion of *luxS* into the FNN ATCC 23726 chromosome

The plasmid pBCG02-luxS7_1 was introduced into electrocompetent FNN ATCC 23726 by electroporation(81). Single-crossover integration of the plasmid into the chromosome via homologous recombination was selected on TSPC agar plates supplemented with 5 µg/mL thiamphenicol. Two to three thiamphenicol-resistant colonies were picked and propagated in TSPC broth containing thiamphenicol, then subcultured once into antibiotic-free TSPC medium to allow recovery of recombinants that had resolved the integrated plasmid via a second crossover event.

The following day, cultures were diluted 10³- and 10⁴-fold in fresh TSPC medium, and 100 µL of each dilution was plated onto TSPC agar containing 2 mM theophylline for counterselection. In the pBCG02 system, theophylline induces expression of the toxin gene *hicA*, thereby eliminating cells that retain the plasmid and enriching for clones that have lost the integrated plasmid after the second recombination event. After 3 days of anaerobic incubation, approximately 20 colonies were randomly selected and streaked onto both non-selective TSPC plates and TSPC plates containing 5 µg/mL thiamphenicol to assess antibiotic sensitivity. All isolates were thiamphenicol-sensitive, consistent with plasmid loss. Ten thiamphenicol-sensitive clones were subsequently analyzed by colony PCR to confirm the expected in-frame insertion of *luxS* at the target chromosomal locus.

### Deletion of *luxS* in FNA strain 7_1

To generate a markerless Δ*luxS* mutant in FNA strain 7_1, the deletion plasmid pBCG10-Δ*luxS* was first introduced into *E. coli* SZU604, which expresses three *F. nucleatum* methyltransferases derived from FNN ATCC 25586(82). Although originally developed to enhance transformation efficiency in strain ATCC 25586, we found that plasmid DNA prepared from SZU604 significantly improved transformation efficiency in strain 7_1 compared with plasmid DNA isolated from standard cloning strains DH5α. Purified plasmid from SZU604 was therefore used for electroporation into electrocompetent FNA 7_1 cells.

In the presence of 2 mM theophylline, riboswitch-controlled *repA* expression allows pBCG10-Δ*luxS* to replicate as a shuttle plasmid under thiamphenicol selection. To obtain chromosomal integrants, transformants were subsequently cultured in medium lacking theophylline while maintaining thiamphenicol selection. Withdrawal of theophylline prevents plasmid replication, and thiamphenicol selection permits survival only of cells in which homologous recombination has integrated the plasmid into the chromosome via a single crossover event.

Integrants were then propagated in non-selective medium to allow recovery of cells in which a second homologous recombination event had occurred, resolving the integrated plasmid. This resolution event can generate either restoration of the wild-type allele or replacement with the in-frame Δ*luxS* allele carried on the plasmid. To eliminate cells that retained plasmid sequences, cultures were plated on medium containing 100 ng/ml anhydrotetracycline (ATc), which induces expression of the MazF toxin encoded on pBCG10. ATc counterselection selectively kills plasmid-retaining cells, enriching for clones that have lost the plasmid backbone following resolution. Surviving colonies were screened for thiamphenicol sensitivity to confirm plasmid loss and subsequently analyzed by colony PCR to identify strains carrying the desired Δ*luxS* mutation.

### Initial attempt to generate an in-frame deletion of *metK*

A previously described *galK*-based counterselection strategy was adapted to generate a non-polar, in-frame *metK* deletion in FNN ATCC 23726(52). The deletion plasmid pCM-galK-Δ*metK* was introduced into competent cells of strain cw1, a Δ*galK* derivative of ATCC 23726, by electroporation. Chromosomal integration of the plasmid via homologous recombination was selected on TSPC agar plates containing 5 µg/mL thiamphenicol at 37°C. Two to three thiamphenicol-resistant colonies were isolated and cultured overnight in TSPC broth without antibiotic selection to allow resolution of the integrated plasmid through a second homologous recombination event. The following day, cultures were diluted 1,000-fold, and 100 µL aliquots were plated onto TSPC agar supplemented with 0.25% 2-deoxy-D-galactose (2-DG). In this system, reconstitution of *galK* renders cells sensitive to 2-DG; therefore, growth on 2-DG selects for cells that have lost the plasmid backbone following the second crossover event. Colonies exhibiting the expected phenotype (2-DG-resistant and thiamphenicol-sensitive) were screened by colony PCR to determine whether the chromosomal *metK* locus had been replaced by the in-frame deletion allele. Approximately 100 candidate colonies were examined; however, all retained the wild-type *metK* allele, and no deletion mutants were recovered. These results strongly suggest that *metK* is essential under the conditions tested.

### Creation of a conditional *metK* deletion mutant

To generate a conditional *metK* mutant, the chromosomal integrant strain harboring pCM-galK-Δ*metK* was complemented in trans with plasmid pSX11E-Pfdx-E-*metK*, which expresses *metK* under the control of a theophylline-inducible promoter. Transformants were selected on TSPC agar plates containing 5 µg/mL thiamphenicol and 1 µg/mL clindamycin to maintain both the integrated deletion construct and the complementing plasmid. A single double-resistant colony was inoculated into TSPC broth supplemented with clindamycin and grown overnight. The culture was then diluted 1:100, and 100-µL aliquots were plated onto TSPC agar containing 2 mM theophylline and clindamycin. Induction with theophylline ensured expression of the plasmid-borne *metK* during subsequent resolution of the chromosomal integrant. After three days of anaerobic incubation at 37°C, colonies were screened for loss of thiamphenicol resistance, indicating excision of the integrated deletion plasmid. Thiamphenicol-sensitive isolates were further analyzed by colony PCR to determine whether the chromosomal *metK* locus had been successfully replaced by the deletion allele.

### AI-2 assay

The AI-2 bioluminescence assay was conducted following established protocols (41, 83). In brief, bacterial cultures in the stationary phase were centrifuged at 12,000 g for 5 minutes to pellet the cells. The resulting supernatant was filtered through a 0.22 μm filter (Millipore, Bedford, MA, USA) to obtain cell-free supernatant (CS) samples. The reporter strain *V. harveyi* BB170 was diluted 1:5000 in fresh autoinducer bioassay medium (consisting of 0.3 M NaCl, 0.05 M MgSO₄, 0.2% Casamino Acids, 10 μM KH₂PO₄, 1 μM L-arginine, 20% glycerol, 0.01 μg/mL riboflavin, and 1 μg/mL thiamine) and cultured at 30°C for 3–4 hours. After incubation, 4.5 ml of the *V. harveyi* culture was mixed with 0.5 ml of the CS sample, and the mixture was incubated at 30°C for 4 hours. After incubation, 200 μL aliquots were transferred to a black flat-bottomed 96-well microplate (Greiner Bio-one: #655096) for bioluminescence measurement using a GloMax® Navigator Microplate Luminometer. Cell-free supernatant from *E. coli* BL-21 was the positive control, while its *luxS* mutant was the negative control. The *V. harveyi* bioassay was performed in triplicate for each sample within a single experiment, and the experiments were repeated three times.

### Cell growth assay

FNN ATCC 23726 wild-type and *luxS* insertion strains, as well as FNA strain 7_1 and its Δ*luxS* mutant, were grown overnight in TSPC broth under anaerobic conditions at 37°C. Overnight cultures were diluted into fresh TSPC medium to an initial OD_600_ of 0.1. Bacterial growth was monitored by measuring optical density at 600 nm (OD_600_) using an Implen NanoPhotometer N60 UV/visible spectrophotometer. For detailed growth curves, OD600 was measured every hour for up to 15 hours. In some experiments, only two times (7 h and 24 h) were measured. Prior to each measurement, culture tubes were vortexed for 10 seconds to ensure a homogeneous cell suspension. The OD values presented are averages of three independent experiments performed in duplicate.

### Biofilm assay

*In vitro* biofilm formation by *F. nucleatum* strains was assessed as previously described (52). Briefly, overnight cultures were diluted 1:100 into fresh TSPC broth (pH adjusted to 8.5), and 3 mL aliquots were dispensed into flat-bottom 12-well plates (Greiner Bio-One). Plates were incubated anaerobically at 37°C for 72 hours to allow biofilm development. Following incubation, planktonic cells were removed, and the biofilms were gently washed twice with sterile water. Plates were air-dried prior to staining with 1% (w/v) crystal violet. Excess dye was removed by washing with water. To quantify biofilm biomass, bound crystal violet was solubilized with 95% ethanol, and absorbance was measured at 580 nm using a Tecan Infinite M1000 microplate reader. All experiments were performed in triplicate and repeated independently three times.

### Gene expression profiling by RNA-seq

For transcriptomic analysis, FNA strain 7_1 and its Δ*luxS* mutant were grown anaerobically in TSPC medium overnight at 37°C. The following day, cultures were diluted 1:20 into fresh TSPC medium and incubated under the same conditions. Cells were harvested at 8 and 12 hours by centrifugation. For FNN ATCC 23726, cultures grew to an OD_600_ of approximately 0.6. The culture was then divided into two aliquots: one was treated with 30 µM synthetic DPD (Omm Scientific), and the other served as an untreated control. After 1.5 hours of incubation under anaerobic conditions, cells were collected by centrifugation. Total RNA was extracted and submitted to the Cancer Genomics Center at The University of Texas Health Science Center at Houston (CPRIT RP240610) for sequencing. RNA quality was assessed using the Agilent RNA 6000 Pico Kit (#5067-1513) on an Agilent 2100 Bioanalyzer (Agilent Technologies, Santa Clara, CA, USA). Samples with an RNA integrity number (RIN) greater than 7 were used for library preparation. Ribosomal RNA was depleted prior to library construction. Strand-specific libraries were prepared using the NEBNext Ultra II Directional RNA Library Prep Kit for Illumina (E7760L, New England Biolabs) together with NEBNext Multiplex Oligos for Illumina (E6609S, New England Biolabs), following the manufacturer’s instructions. Final library quality was evaluated using the Agilent High Sensitivity DNA Kit (5067–4626) on the Agilent 2100 Bioanalyzer, and library concentrations were determined by quantitative PCR using the Collibri Library Quantification Kit (A38524500, Thermo Fisher Scientific). Libraries were pooled at equimolar concentrations and subjected to paired-end 150-bp sequencing on an Illumina NovaSeq platform (Illumina, USA). Raw paired-end reads were processed and aligned to the reference genome, and differential gene expression analysis was performed using DESeq2. Downstream analyses and data visualization were conducted using R/Bioconductor packages. Genes with an absolute log₂(fold change) ≥ 1.0 and a p-value ≤ 0.05 were considered differentially expressed under the tested conditions.

### Transmission Electron Microscopy

The conditional *metK* mutant was grown overnight in TSPC medium supplemented with 2 mM theophylline. The following day, cultures were washed twice with fresh TSPC medium and diluted 1:20 into fresh medium containing either 2 mM theophylline or no theophylline (MetK depletion). After 18 hours of anaerobic incubation, cells were harvested by centrifugation, washed with 100 mM NaCl, and resuspended in the same solution.

A 10-µL aliquot of each bacterial suspension was applied onto carbon-coated nickel grids and allowed to adsorb. Grids were negatively stained with 0.1% (w/v) uranyl acetate and briefly rinsed with sterile water. Samples were examined using a JEOL JEM-1400 transmission electron microscope.

## Data Availability

The RNA-seq raw data were deposited in the NCBI Sequence Read Archive (SRA) under the accession numbers PRJNA1275848 and PRJNA1276057. Plasmid pBCG10 (232323) has been deposited with Addgene.

## AUTHOR CONTRIBUTIONS

C.W. and B. G conceived and designed all experiments. B.G., S.X, T. T, and C.W. performed all experiments. B.G. and C.W. analyzed data. C. W. and B.G wrote the manuscript with contributions and approval from all authors.

## ACKNOWLEDGMENTS

This work was supported by NIDCR grants DE030895 and DE034542 awarded to C.W. We thank the technical support (RNA-seq) from the Cancer Prevention and Research Institute of Texas (CPRITPR240610). We are grateful to Dr. Dnaiel Slade (Virginia Tech) for providing the FNA strain 7_1 and Dr. Xiangan Han (Shanghai Veterinary Research Institute, Chinese Academy of Agricultural Sciences) for providing the *E.coli* L-21 (DE3) Δ*luxS* strain.

